# Whole-genome sequencing of Vero E6 (C1008) and comparative analysis of four Vero cell sublines

**DOI:** 10.1101/2021.10.26.466002

**Authors:** Kazuhiro Konishi, Toshiyuki Yamaji, Chisato Sakuma, Fumio Kasai, Toshinori Endo, Arihiro Kohara, Kentaro Hanada, Naoki Osada

## Abstract

The Vero cell line is an immortalized cell line established from kidney epithelial cells of the African green monkey. A variety of sublines have been established from the original cell line, which display different characteristics. In this study, we determined the whole-genome sequence of Vero E6 (C1008) and performed comparative analysis among Vero JCRB 0111, Vero CCL-81, Vero 76 and Vero E6. Analysis of the copy number changes and loss of heterozygosity revealed that all sublines share a large deletion and loss of heterozygosity on chromosome 12, which harbors type I interferon and *CDKN2* gene clusters. We identified a substantial number of genetic differences among the sublines including single nucleotide variants, indels, and copy number variations. The spectrum of single nucleotide variants indicated a close genetic relationship between Vero JCRB0111 and Vero CCL-81, and between Vero 76 and Vero E6, and a considerable genetic gap between the former two and the latter two lines. In contrast, we confirmed the pattern of genomic integration sites of simian endogenous retroviral sequences, which was consistent among the sublines. We identified subline-specific/enriched loss of function and missense variants, which potentially contribute to the differences in response to viral infection among the Vero sublines. In particular, we focused on Vero E6-specific/enriched variants and identified four genes (IL1RAP, TRIM25, RB1CC1, and ATG2A) that contained missense variants specific or enriched in Vero E6. In addition, we found that V739I variants of ACE2, which functions as the receptor for SARS-CoV-2, were heterozygous in Vero JCRB0111, Vero CCL-81, and Vero 76; however, Vero E6 contained the allele with isoleucine, resulting from the loss of one of the X chromosomes.

## Introduction

Cell lines established from mammalian tissues are often used for virus isolation and culture as well as for vaccine production. One of the cell lines frequently used for these purposes is the Vero cell line, which is an immortalized cell line established from kidney epithelial cells of an African green monkey (*Chlorocebus sabaeus*, AGM) by Yoshihiro Yasumura at Chiba University in 1962. After the cell lines were established, they were brought to the National Institute of Allergy and Infectious Diseases in the US, the American Type Culture Collection (ATCC), and the Japanese Collection of Research Bioresources (JCRB) cell bank. Various sublines have been established through passaging (Earley and Johnson 1988; Mizusawa, Simizu, and Terasima 1988; Terasima, Yasukawa, and Simizu 1988), which have slightly different properties from one another. For example, Vero CCL-81 (ATCC CCL-81) is known to be more likely to propagate Japanese encephalitis virus, and Vero E6 (C1008) is more likely to propagate SARS-CoV-2 compared with the other sublines (Harcourt et al. 2020; Matsuyama et al. 2020). However, the genetic factors that contribute to these phenotypic differences are largely unknown.

A whole-genome sequencing analysis of one of the sublines, Vero JCRB0111 was performed by Osada et al. (Osada et al. 2014) and approximate 9Mbp homozygous deletion on chromosome 12 was identified. The deletion was found to contain a cluster of type 1 interferon (*IFN*-1) genes that act as viral suppressors, as well as *CDKN2A* and *CDKN2B*, which are involved in the cell cycle. These results indicate potential factors that resulted in the immortalization of Vero cells and may partly explain why viruses can readily multiply in these cells. Furthermore, whole-genome resequencing analysis of additional sublines, Vero CCL-81 and Vero 76 was performed by Sakuma et al. (Sakuma et al. 2018), in which they identified numerous genetic variations among the sublines as well as integrations of Simian endogenous retroviral sequences (SERVs); however, the effects of the variants at the nucleotide level were not evaluated. More recently, Sene et al. reported the haplotype-resolved genome assembly of the Vero CCL-81 subline (Sène et al. 2021). Hundreds of protein-coding genes, including *ACE2*, were identified as losing function. Although the intrinsic function of ACE2 is an angiotensin-converting, it is also known as the host cell receptor for SARS-CoV and SARS-CoV-2. Overall, these studies have provided information for quality control and have produced novel engineered sublines, which will accelerate the development of vaccine manufacturing platforms.

In the present study, we performed genome sequencing of Vero E6, which was established from a clone isolated from Vero 76, and conducted a comparative genome analysis among four different Vero cell sublines, including Vero JCRB0111, Vero CCL-81, Vero 76, and Vero E6. The inferred passaging history of these sublines is shown in Figure 1. The identification of genetic variants specific or enriched in particular sublines will contribute to the elucidation of factors responsible for the phenotypic differences among the sublines.

**Figure 1.**
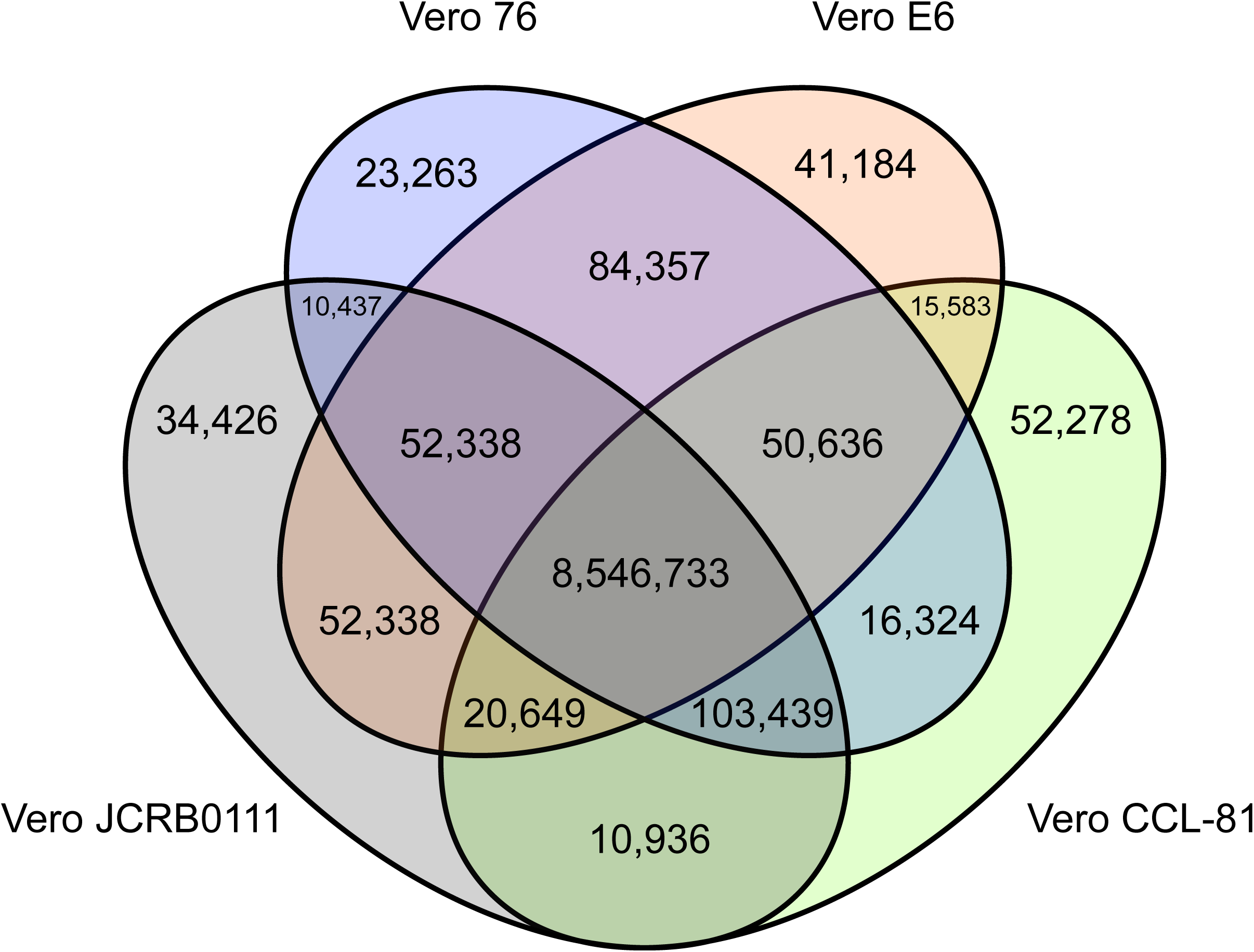
History of Vero cell lines. Figure is adopted from Sakuma et al. (2018) under a Creative Commons Attribution 4.0 International license.

## Materials and Methods

### Data preparation

Vero E6 cells were obtained from the JCRB cell bank. The short-read sequences of Vero JCRB0111, Vero CCL-81, and Vero 76 were downloaded from a public database (DDBJ: PRJDB2865). Paired-end sequences of Vero E6 were determined using an Illumina HiSeq 2500. The reads were mapped to the AGM reference genome (NCBI: GCF_000409795.2) using the BWA-mem algorithm (Li and Durbin 2009). The single nucleotide variants (SNVs) and short indels (<50 bp) were called using VarScan 2 software (Koboldt et al. 2012). The thresholds for SNV detection were: ≥13 coverage, ≥2 mutation count, ≥15 average base quality. The minimum coverage was selected to cover at least 95% of the genome for each sample. The strand filter was applied to exclude the sites where ≥90% of the reads mapped to one strand. Only bi-allelic SNVs were considered in this study. Manta was used to identify large-scale structural variations (Chen et al. 2015).

The GTF format file downloaded from the Ensembl database was used for gene annotation (Chlorocebus_sabaeus.ChlSab1.1.86.gtf) and snpEFF software was used for annotating the effect of each genetic variation (Cingolani et al. 2012). The assignment for the gene function categories was performed using DAVID (Jiao et al. 2012). For evaluating the impact of missense variants, we used PROVEAN, SIFT, and PANTHER-PSEP (Choi et al. 2012; Choi and Chan 2015; Tang and Thomas 2016). The RNA-seq experiment data of Vero E6 TMPRSS2+ cells were downloaded from the public database (SRR13091741–SRR13091746) (Zhang et al. 2021). The RNA-seq reads were mapped to the reference genome using HISAT2 using the default parameters (Kim et al. 2019).

We defined subline-specific/enriched SNVs as variants with a significantly higher frequency compared with the other sublines. For each SNV, we counted the number of reads supporting the reference and alternative alleles and performed Tukey’s test (Tukey 1949). SNVs with *P*<0.05 were considered subline-specific/enriched SNVs after controlling for multiple-testing using the Benjamini-Hochberg method (Benjamini and Hochberg 1995).

### Copy number variations and loss of heterozygosity

We used Control-FREEC software to identify copy number variations (CNVs) and loss of heterozygosity (LOH) (Boeva et al. 2012). A window size of 50 kbp and a step size of 10 kbp were selected. To quantitate the genetic differentiation between sublines, we computed *f*_2_ statistics between all pairwise sublines. *F*_2_ was calculated as the sum of the squares of allele frequency differences between two sublines, divided by the total number of SNVs. The *f*_2_ values were used as a genetic distance between two sublines and a neighbor-joining tree was constructed using MEGA X (Knyaz et al. 2018).

### Genomic PCR for SVL27b, a SERV-variation locus on chromosome 27

PCR was performed as described previously (Sakuma et al. 2018). Genomic DNA (30 ng) was used as the template. The primers specified below were synthesized (Eurofins Genomics Inc., Tokyo, Japan): Fw1: 5′-GGAACACCTGAAGATCTATGTGTCTA-3′, Rv1: 5′-ATCAAATTCCTCTCTTCACATCTTCT-3′, Fw2: 5′-GGACATATTGTTATAAAAGTTCATGG-3′, Rv2: 5′-GAAACTATACCTATGATTTTGCCATAG-3′

## Results and discussion

### Genomic features of the Vero sublines

We obtained short reads from Vero E6 and mapped to the reference genome of AGM. To compare the genomic features of the Vero sublines, we reanalyzed previously published genomic data (Vero JCRB0111, Vero CCL-81, and Vero 76) and the newly obtained genome sequence of Vero E6 cells. In total, we obtained 74-, 32-, 43-, and 78-fold coverage of the Vero JCRB00111, Vero CCL-81, Vero 76, and Vero E6 genomes, respectively. A 20% frequency cut-off yielded 8,867,211 SNVs, 453,797 short insertions, and 547,598 short deletions. As expected, 94% of these variants were shared among all sublines. The number of SNVs shared among sublines is presented in Figure 2.

**Figure 2.**
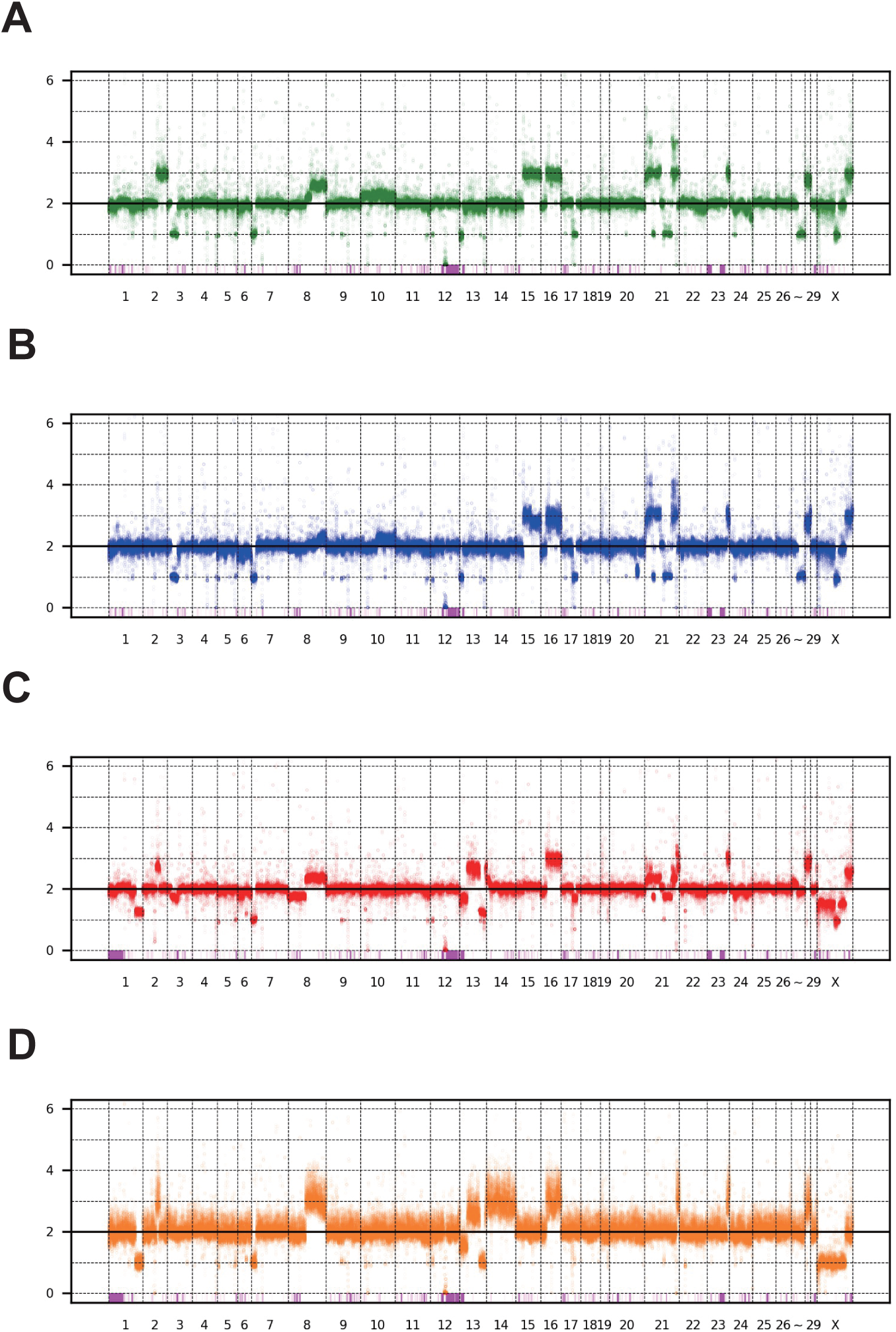
Venn diagram of SNVs shared among the four different Vero sublines. The SNVs with frequency >0.2 are labeled as present in the subline.

The patterns of CNV and LOH are shown in Figure 3. Overall, the patterns appeared similar among the four sublines, including the 9 Mbp deletion on chromosome 12. However, there were several marked differences; for example, a large part of chromosome 15 had three copies in Vero JCRB0111 and Vero CCL-81 but the region was normal with respect to copy number in Vero 76 and Vero E6. Interestingly, chromosome 21 is highly rearranged in Vero JCRB0111, Vero CCL-81, and Vero 76, but the copy number of chromosome 21 in Vero E6 is normal, except for the distal end. Vero E6 also showed monosomy for the X chromosome. This feature is partially observed in Vero 76, showing a mosaic copy number for the X chromosome. This indicates that the Vero 76 cell line is composed of a heterogeneous cell population, which may be distinguished by one or two copies of the X chromosome.

**Figure 3.**
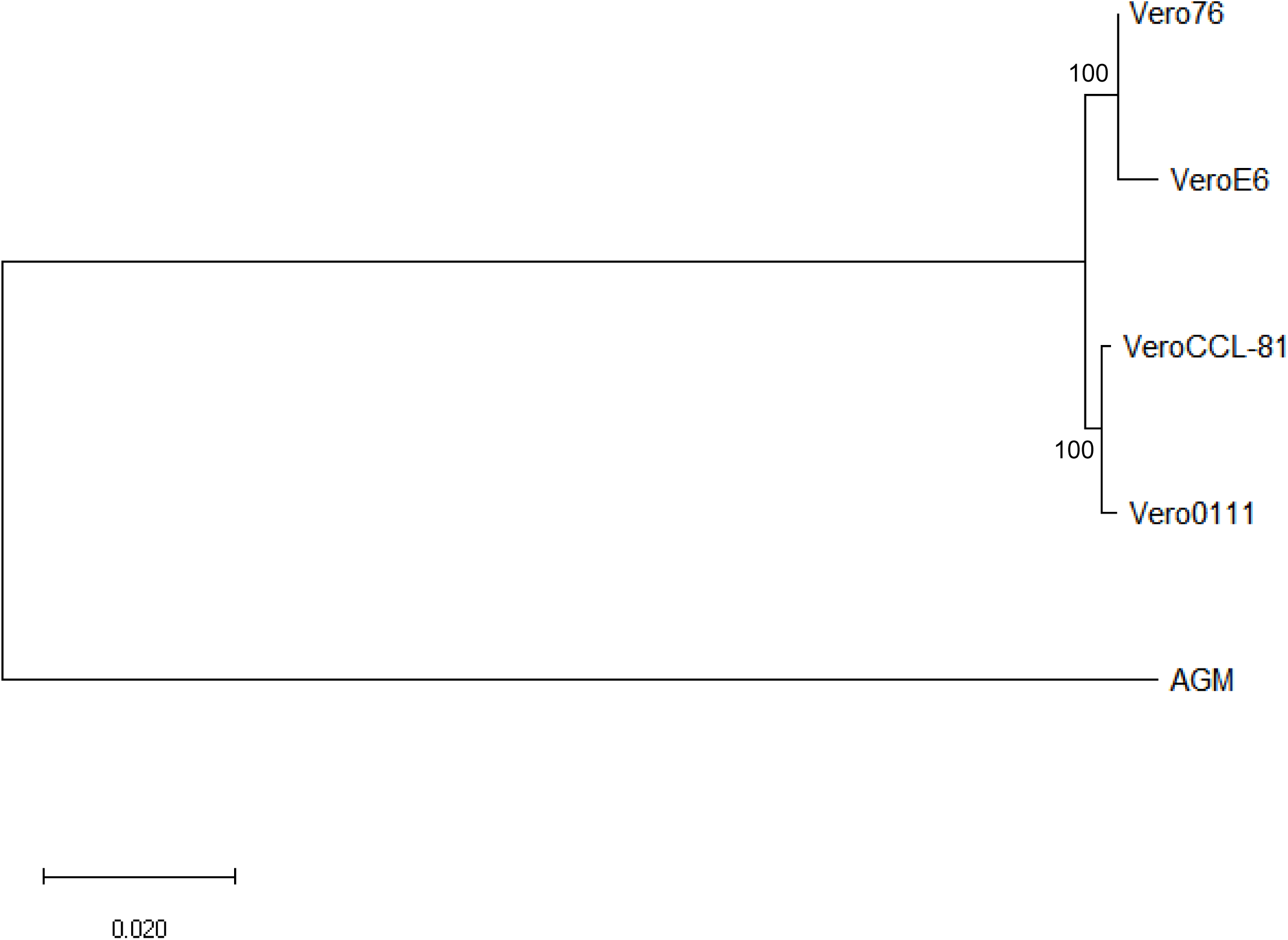
Copy number variation of four the Vero sublines. The dots represent the estimated copy number in 50 kb-length windows. X- and y-axes denote chromosomal coordinates and estimated copy numbers, respectively. The chromosome number is labeled under the panel. The pink boxes represent the regions of LOH.

### Genetic relatedness among sublines

Next, we evaluated the genetic relationship among the four sublines. We used *f*_2_ statistics to measure the genetic distance between the sublines. *F*_2_ statistics are population genetics based statistics that measure the difference of the frequency of variants between two cell populations (Patterson et al. 2012). The neighbor-joining tree reconstructed using *f*_2_ statistics is shown in Figure 4. The tree indicates the sister relationship between Vero JCRB0111 and Vero CCL-81, and between Vero 76 and Vero E6 with 100% bootstrap support. The reconstructed tree was incongruent with the inferred history of the Vero cell lineages (Figure 1). This may have resulted from rapid turnover of cell karyotypes within the sublines. The frequency of clonal cell lineages may change over time because of the difference in the survival/proliferation rate or by other unknown mechanisms. The results suggest that different karyotypes co-existed before the split of Vero JCRB0111 and Vero CCL-81, which increased in frequency independently in Vero JCRB0111-CCL81 and Vero 76-E6. Further studies of karyotyping and single-cell genomic sequencing will reveal the dynamics of the cell populations during subline divergence.

**Figure 4.**
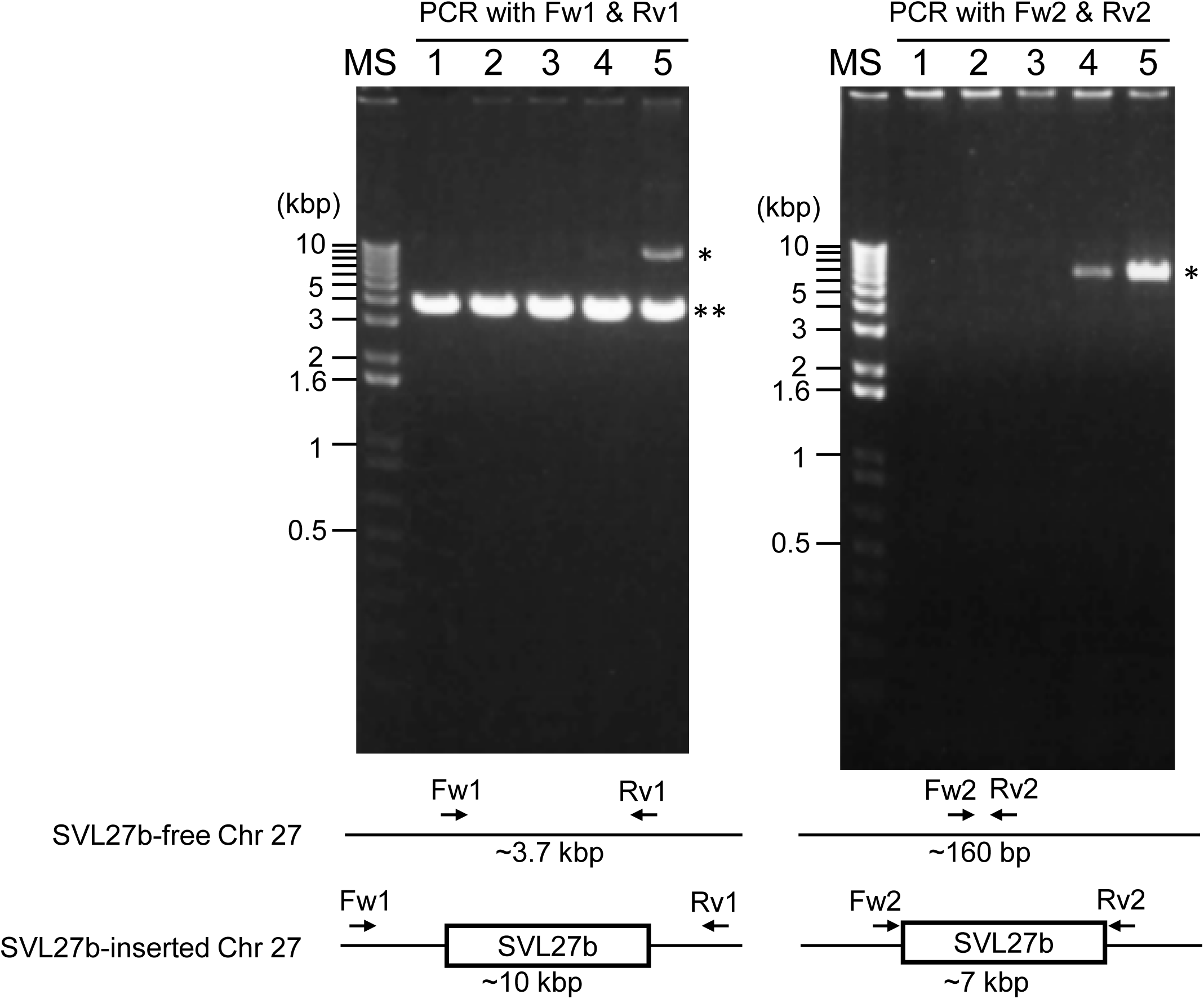
Cladogram of the four Vero sublines. The genetic distances were measured using *f*_2_ statistics and the neighbor-joining method was used for the tree reconstruction. Bootstrap % values are shown on the blanches.

### Survey of endogenous retroviral sequences

The study of Sakuma et al. (Sakuma et al. 2018) revealed many SERV sequences that are present in the genomes of Vero JCRB0111, Vero CCL-81, and Vero 76, however, the pattern of insertions was perfectly consistent among the three sublines except for one insertion, which was referred to as SVL27b. SVL27b is present in the genome of Vero 76 but absent in the genomes of Vero JCRB0111 and Vero CCL-81 (Sakuma et al. 2018). Considering that the African green monkey individual sequenced for draft genome harbored the insertion as a heterozygous state, Sakuma et al. (2018) concluded that the insertion was lost in Vero JCRB0111 and CCL-81 but retained in Vero 76. In this study, we investigated the pattern of SERV insertions in Vero E6, following the method of Sakuma et al. (2018) and confirmed that the integration pattern was the same as Vero 76, except for the one on the X chromosome. The SERV insertion around chrX:47012301-47012900 is heterozygous in Vero JCRB0111, Vero CCL-81, and Vero 76, but absent in Vero E6. The result is consistent with the CNV analysis, which showed that Vero E6 lost one of the two X chromosomes in almost every cell.

Whole-genome sequence analysis also showed that SVL27b is present in the Vero E6 genome, which is consistent with the idea that Vero E6 is a clonal derivative from Vero 76 cells (Earley and Johnson 1988). This was verified by genomic PCR experiments as shown in Figure 5. When a PCR primer set matching a region ∼2-kbp away from the SVL27b inserted site, an approximate 3.7 kb DNA fragment corresponding to AGM chromosome 27, without the SERV insertion at this position, was amplified from all four Vero cell lines as well as the AGM control (Figure 5). This indicates that the Vero cell lines have an allele(s) that do not contain SVL27b. Notably, a DNA fragment with an ∼7 kb SVL27b insertion was amplified from Vero E6 (Figure 5). When a PCR primer was designed proximal to the SVL27b inserted site, a DNA fragment with the SVL27b insert was amplified from Vero 76 and Vero E6 cells, but not from the others (Figure 5). The DNA band amplified from Vero E6 was much denser compared with of Vero 76 (Figure 5). It should also be pointed out that, under the latter PCR conditions, amplified DNA without the SVL27b insertion was too short to be visible using agarose gel electrophoresis. Together with the results of the previous study (Sakuma et al. 2018), these results strongly suggest that Vero 76 cells are a mixture of cell types with and without the SVL27b insertion and Vero E6 is a Vero 76-derived clone which stably has SVL27b. Furthermore, these results indicate that SVL27b is a good genomic marker to distinguish the Vero 76-Vero E6 lineage from the Vero CCL-81 and JCBR0111 lineages (Fig. 1).

**Figure 5.**
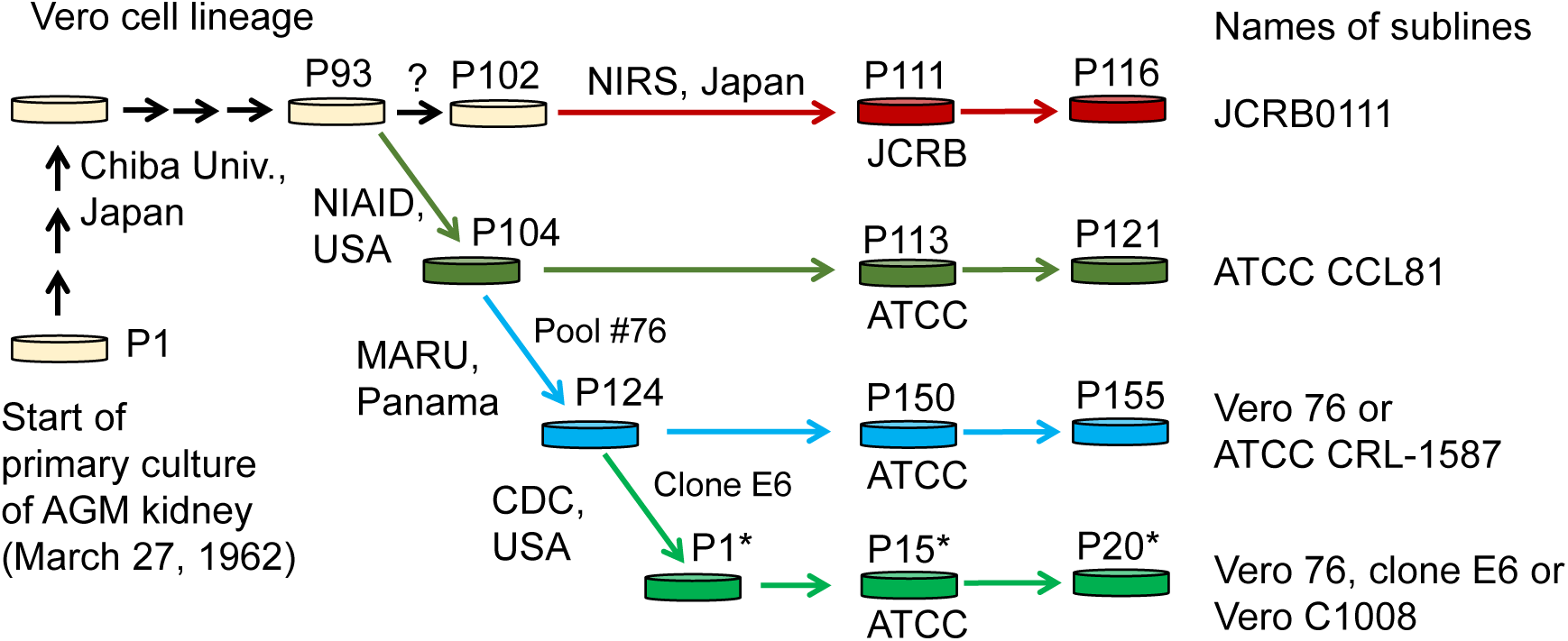
Using genomic DNA prepared from various cells as a template, DNA fragments were amplified by PCR with two different combinations of primers. Sequences of the primers are specified in the Materials and Methods. 1) AGM lymphocytes; 2) Vero JCRB0111; 3) Vero ATCC CCL81; 4) Vero 76; 5) Vero E6 (C1008). *, fragment containing SERV; **, fragment not containing SERV.

### Identification of subline-specific/enriched variants

We next looked into the genetic variants specifically observed or exhibiting a high frequency in each subline. We defined a subline-specific/enriched variant as a variant with higher frequency compared with the other sublines with statistical significance (see method). In total, 22, 107, 28,985, 6705, 122,198 variants were labeled as specific/enriched in Vero JCRB0111, Vero CCL-81, Vero 76, and Vero E6, respectively. Vero E6 showed the highest number of subline-specific/enriched variants. For the genes with missense SNVs, we also checked the RNA-seq expression data and retained the genes only if the mRNA with variant alleles was actually expressed in Vero E6 TMPRSS2+ cells. In total, 66, 48, 12, and 272 subline-specific/enriched missense SNVs were found on 47, 35, 12, and 188 protein-coding genes in Vero JCRB0111, Vero CCL-81, Vero 76, and Vero E6 cells, respectively (Table 1, Supplementary Table 1). In addition, we identified 4, 8, 2, and 29 genes that harbor subline-specific loss-of-function (LOF) variants resulting from SNVs and short indels in Vero JCRB0111, Vero CCL-81, Vero 76, and Vero E6 cells, respectively (Supplementary Table 2). The analysis of large structural variants using Manta software also revealed 465, 12, 52, and 682 genes with LOH variants because of structural variations (Supplementary Table 3).

**Table 1.** Subline-specific/enriched missense variants.

|  | Vero JCRB0111 | Vero CCL-81 | Vero 76 | Vero E6 |
| --- | --- | --- | --- | --- |
| Missense SNV | 66 | 48 | 12 | 272 |
| # of affected genes | 47 | 35 | 12 | 188 |

**Table 2.** Subline-specific LOF variants in genes related to viral infection/proliferation.

|  | Vero<br>JCRB0111 sp. | Vero<br>CCL-81 sp. | Vero 76 sp. | Vero E6 sp. |
| --- | --- | --- | --- | --- |
| SNV | 2 | 3 | 0 | 7 (1) |
| Insertion | 0 | 0 | 2 | 4 |
| Deletion | 2 | 5 | 0 | 18 (2) |
| # of affected<br>genes | 4 | 8 | 1 | 29 |

\*The numbers of homozygous variants are shown in the parentheses.

We further surveyed genes related to autophagy, apoptosis, antiviral activity, or innate immune response and harbored subline-specific/enriched missense or LOH variants in a single subline. For this purpose, we excluded the structural variants identified by Manta, because candidate variants could contain a nonnegligible amount of false-positive variants and required additional experimental validations (Kawamoto et al. 2020). In total, we identified eight genes with subline-specific/enriched missense SNVs (Table 3). On the other hand, none of the genes with subline-specific/enriched LOH variants have functions related to the above functional categories.

**Table 3.**
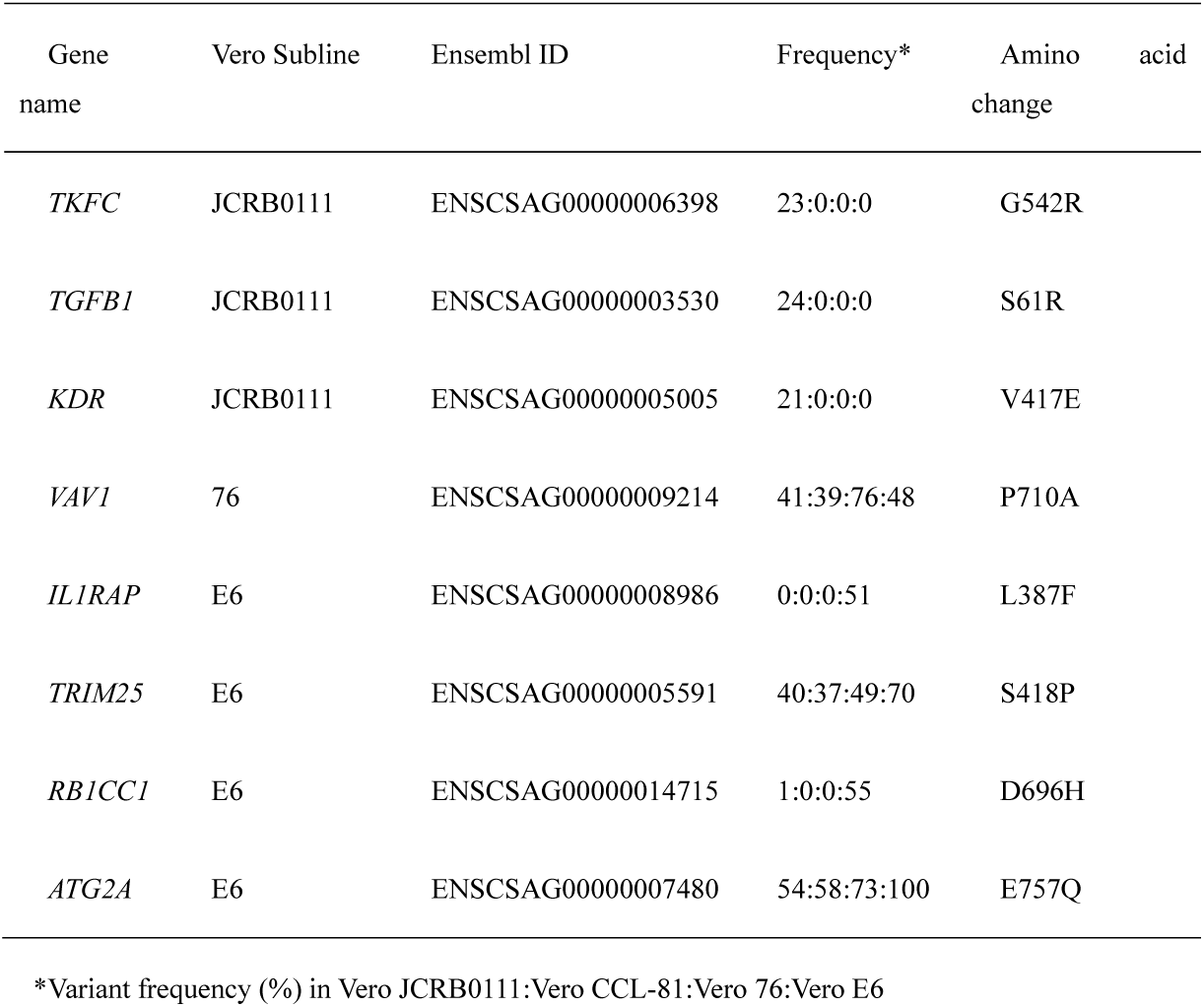
Subline-specific/enriched missense SNVs in genes related to viral infection/proliferation.

| Gene name | Vero Subline | Ensembl ID | Frequency* | Amino acid change |
| --- | --- | --- | --- | --- |
| *TKFC* | JCRB0111 | ENSCSAG00000006398 | 23:0:0:0 | G542R |
| *TGFB1* | JCRB0111 | ENSCSAG00000003530 | 24:0:0:0 | S61R |
| *KDR* | JCRB0111 | ENSCSAG00000005005 | 21:0:0:0 | V417E |
| *VAV1* | 76 | ENSCSAG00000009214 | 41:39:76:48 | P710A |
| *IL1RAP* | E6 | ENSCSAG00000008986 | 0:0:0:51 | L387F |
| *TRIM25* | E6 | ENSCSAG00000005591 | 40:37:49:70 | S418P |
| *RB1CC1* | E6 | ENSCSAG00000014715 | 1:0:0:55 | D696H |
| *ATG2A* | E6 | ENSCSAG00000007480 | 54:58:73:100 | E757Q |

\*Variant frequency (%) in Vero JCRB0111:Vero CCL-81:Vero 76:Vero E6

### Variants in Vero E6

W focused on genetic variants specific/enriched in Vero E6, which efficiently propagate SARS-CoV-2. Missense variants in *IL1RAP* (L387F) and *BR1CC1* (D696H) were almost exclusively found in Vero E6 in a heterozygous state, whereas those in *TRIM25* (S418P) and *ATG2A* (E757Q) had a statistically high frequency in Vero E6.

IL1RAP is an auxiliary receptor of IL1R1, a receptor for IL-1α and IL-1β. Virus-induced cell death releases IL-1α in the cytoplasm (Malik and Kanneganti 2018), and the released IL-1α binds to IL1R1 on the plasma membrane. The intracellular toll IL-1 receptor (TIR) domain of IL1R1 binds to IL1RAP, leading to the activation of key transcription factors and kinases associated with the inflammatory and immune response, such as NF-κB, AP1, JNK, MAPK, and ERK (Mantovani et al. 2019). The leucine residue at 387 of the IL1RAP protein is highly conserved among mammals and prediction programs reported that the variant, leucine to phenylalanine, would be deleterious. Therefore, the L387F amino acid change in IL1RAP may disrupt the downstream cascade, suppressing the inflammatory and immune response and facilitating the increase in SARS-CoV-2 infection.

RB1CC1 constitutes a part of the ULK1 complex, which is required for the initiation of autophagy (Bello-Perez et al. 2020). The ULK1 complex phosphorylates a part of the PI3KC3 complex, which triggers the formation of a phagophore, and known as the sequestration membrane (Dikic and Elazar 2018). Phagophores envelop viruses and virus-derived antigens to form autophagosomes, and the fusion of lysosomes and autophagosomes results in the formation of autolysosomes and degradation of their contents (Ahmad, Mostowy, and Sancho-Shimizu 2018). Autophagosomes fuse with endosomes to form amphisomes, and amphisomes fuse with lysosomes to form autolysosomes (Zhao and Zhang 2018). The endosomes may contain SARS-CoV-2 that has entered via endocytosis (Bian and Li 2021). Previous studies have shown that ORF3a of SARS-COV-2 inhibits two pathways that form autolysosomes, suggesting that it prevents itself from being degraded by inhibiting autophagy (Miao et al. 2021). The prediction programs suggested that the D696H variant would be deleterious and deteriorate its function. Therefore, the occurrence of non-synonymous variants in RB1CC1 may have inhibited the initiation of autophagy and suppressed the degradation of SARS-CoV-2 by autophagy.

TRIM25 is a ubiquitin ligase that regulates RIG-1 (DDX58), which detects viral RNA and triggers the innate immune responses (Zeng et al. 2010). TRIM25 causes polyubiquitination in a region of RIG-1 called CARD, which activates RIG-1 (Gack et al. 2007). Once activated, RIG-1 recognizes viral RNA, it triggers the activation of NF-κB, IRF3, IRF7, which induces the expression of the antiviral protein Viperin (Schneider, Chevillotte, and Rice 2014). Viperin catalyzes the production of ddhCTP, which interferes with RNA polymerase (RdRp), to promote the degradation of viral proteins, and interferes with the transport of viral proteins (Rivera-Serrano et al. 2020). However, the amino acid site 418 in TRIM25 is relatively variable among other vertebrate orthologs and PROVEN software predicted that the variant of S418P would be functionally neutral.

ATG2A is a protein required for phagophore membrane expansion (Kotani et al. 2018). Previous studies have shown that silencing ATG2A results in the accumulation of unclosed autophagosomes (Velikkakath et al. 2012). The E757Q change in ATG2A was predicted as functionally tolerated by PROVEN and SFIT but possibly damaging by PANTHER score.

We also investigated the genetic variants in the *ACE2* gene of Vero E6. Sene et al. reported that the *ACE2* gene in Vero CCL-81 potentially harbors structural variants causing LOF. However, using the publicly available RNA-seq data (Zhang et al. 2021), we confirmed that *ACE2* mRNA is properly expressed in the Vero E6 TMPRSS2+ cells both in the presence and absence of SARS-CoV-2. This indicates that *ACE2* is expressed in Vero cells, although the angiotensin-converting enzymatic activity of the gene product is lost as shown by Sene et al. (2021). We identified one missense SNV that showed marked differences between Vero E6 and the other sublines. The missense variants of V739I in *ACE2* were heterozygous in Vero JCRB0111, Vero CCL-81, and Vero 76, but Vero E6 harbors only the isoleucine allele. The difference results from the fact that Vero E6 exhibits monosomy X.

## Conclusion

In this study, we determined the whole-genome sequence of Vero E6, which has been widely used for the study of SARS-CoV-2 and performed comparative genomics on four different sublines derived from Vero cells. Genomic resources for the Vero cell lines will benefit quality control of vaccine-producing cell substrates. In addition, finding candidates genes contributing to the different phenotypes of the cell lines will facilitate the identification of mechanisms of viral proliferation and the development of effective and safe substrates for vaccine production. The primary goal of this study was to present a whole-genome sequence of Vero E6 as research resources and catalog a list of candidate variants that potentially affect the phenotypic differences among the Vero sublines. The validation of each effect using additional sequencing and experiments will be necessary, although it is beyond the scope of the present study. Despite of these limitations, we provide a list of genetic differences among the four sublines, as well as variants specific or enriched in particular sublines, which represent a valuable resource for quality control of cell lines and understanding the mechanisms of viral proliferation.

## Conflict of Interest

The authors declare that the research was conducted in the absence of any commercial or financial relationships that could be construed as a potential conflict of interest.

## Author Contributions

FK, AK, KH, TE, and NO: study design. TY and CS: performing experiments. KK and NO: data analysis. KH, FK, and NO: writing manuscript. All authors contributed to the article and approved the submitted version.

## Funding

This work was supported by the MEXT KAKENHI (No. JP21H02630) and AMED-CREST (No. JP20gm0910005j0006) to KH

## Data Availability

The short reads were deposited to the DDBJ DRA database under the accession number DRX311507.

## Supplementary Tables

Supplementary Table 1

List of all missense SNVs specific to sublines

Supplementary Table 2

List of all genes with LOF variants specific to sublines

Supplementary Table 3

List of genes with structural variants specific to sublines

## Notes

### Competing Interest Statement

The authors have declared no competing interest.

## References

Ahmad, L., S. Mostowy, and V. Sancho-Shimizu. 2018. Autophagy-Virus Interplay: From Cell Biology to Human Disease. Frontiers in Cell and Developmental Biology 6.

Bello-Perez, M., I. Sola, B. Novoa, D. J. Klionsky, and A. Falco. 2020. Canonical and Noncanonical Autophagy as Potential Targets for COVID-19. Cells 9:1619.

Benjamini, Y., and Y. Hochberg. 1995. Controlling the False Discovery Rate: A Practical and Powerful Approach to Multiple Testing. Journal of the Royal Statistical Society. Series B (Methodological) 57:289–300.

Bian, J., and Z. Li. 2021. Angiotensin-converting enzyme 2 (ACE2): SARS-CoV-2 receptor and RAS modulator. Acta Pharmaceutica Sinica B 11:1–12.

Boeva, V., T. Popova, K. Bleakley, P. Chiche, J. Cappo, G. Schleiermacher, I. Janoueix-Lerosey, O. Delattre, and E. Barillot. 2012. Control-FREEC: a tool for assessing copy number and allelic content using next-generation sequencing data. Bioinformatics 28:423–425.

Chen, X., O. Schulz-Trieglaff, R. Shaw, B. Barnes, F. Schlesinger, M. Källberg, A. J. Cox, S. Kruglyak, and C. T. Saunders. 2015. Manta: rapid detection of structural variants and indels for germline and cancer sequencing applications. Bioinformatics 32:1220–1222.

Choi, Y., and A. P. Chan. 2015. PROVEAN web server: a tool to predict the functional effect of amino acid substitutions and indels. Bioinformatics 31:2745–2747.

Choi, Y., G. E. Sims, S. Murphy, J. R. Miller, and A. P. Chan. 2012. Predicting the Functional Effect of Amino Acid Substitutions and Indels. PLoS ONE 7:e46688.

Cingolani, P., A. Platts, L. L. Wang, M. Coon, T. Nguyen, L. Wang, S. J. Land, X. Lu, and D. M. Ruden. 2012. A program for annotating and predicting the effects of single nucleotide polymorphisms, SnpEff. Fly (Austin) 6:80–92.

Dikic, I., and Z. Elazar. 2018. Mechanism and medical implications of mammalian autophagy. Nature Reviews Molecular Cell Biology 19:349–364.

Earley, E., and K. Johnson. 1988. The lineage of the Vero, Vero 76 and its clone C1008 in the United States. Vero cells: origin, properties and biomedical applications. Chiba University, Tokyo, Japan:26–29.

Gack, M. U., Y. C. Shin, C.-H. Joo et al. 2007. TRIM25 RING-finger E3 ubiquitin ligase is essential for RIG-I-mediated antiviral activity. Nature 446:916–920.

Harcourt, J., A. Tamin, X. Lu et al. 2020. Severe Acute Respiratory Syndrome Coronavirus 2 from Patient with Coronavirus Disease, United States. Emerging Infectious Disease journal 26:1266.

Jiao, X., B. T. Sherman, D. W. Huang, R. Stephens, M. W. Baseler, H. C. Lane, and R. A. Lempicki. 2012. DAVID-WS: a stateful web service to facilitate gene/protein list analysis. Bioinformatics 28:1805–1806.

Kawamoto, M., T. Yamaji, K. Saito, K. Satomura, T. Endo, M. Fukasawa, K. Hanada, and N. Osada. 2020. Identification of characteristic genomic markers in human hepatoma Huh7 and Huh7.5.1-8 cell lines. bioRxiv:2020.2002.2017.953281.

Kim, D., J. M. Paggi, C. Park, C. Bennett, and S. L. Salzberg. 2019. Graph-based genome alignment and genotyping with HISAT2 and HISAT-genotype. Nat. Biotechnol. 37:907–915.

Knyaz, C., G. Stecher, M. Li, S. Kumar, and K. Tamura. 2018. MEGA X: Molecular Evolutionary Genetics Analysis across Computing Platforms. Mol. Biol. Evol. 35:1547–1549.

Koboldt, D. C., Q. Zhang, D. E. Larson, D. Shen, M. D. Mclellan, L. Lin, C. A. Miller, E. R. Mardis, L. Ding, and R. K. Wilson. 2012. VarScan 2: Somatic mutation and copy number alteration discovery in cancer by exome sequencing. Genome Res. 22:568–576.

Kotani, T., H. Kirisako, M. Koizumi, Y. Ohsumi, and H. Nakatogawa. 2018. The Atg2-Atg18 complex tethers pre-autophagosomal membranes to the endoplasmic reticulum for autophagosome formation. Proceedings of the National Academy of Sciences 115:10363–10368.

Li, H., and R. Durbin. 2009. Fast and accurate short read alignment with Burrows–Wheeler transform. Bioinformatics 25:1754–1760.

Malik, A., and T.-D. Kanneganti. 2018. Function and regulation of IL-1α in inflammatory diseases and cancer. Immunol. Rev. 281:124–137.

Mantovani, A., C. A. Dinarello, M. Molgora, and C. Garlanda. 2019. Interleukin-1 and Related Cytokines in the Regulation of Inflammation and Immunity. Immunity 50:778–795.

Matsuyama, S., N. Nao, K. Shirato et al. 2020. Enhanced isolation of SARS-CoV-2 by TMPRSS2-expressing cells. Proceedings of the National Academy of Sciences 117:7001–7003.

Miao, G., H. Zhao, Y. Li, M. Ji, Y. Chen, Y. Shi, Y. Bi, P. Wang, and H. Zhang. 2021. ORF3a of the COVID-19 virus SARS-CoV-2 blocks HOPS complex-mediated assembly of the SNARE complex required for autolysosome formation. Dev. Cell 56:427–442.e425.

Mizusawa, H., B. Simizu, and T. Terasima. 1988. Cell line Vero deposited to Japanese Cancer Research Resources Bank. VERO cells: origin, properties and biomedical applications. Department of Microbiology School of Medicine Chiba University, Chiba, Japan:24–25.

Osada, N., A. Kohara, T. Yamaji, N. Hirayama, F. Kasai, T. Sekizuka, M. Kuroda, and K. Hanada. 2014. The Genome Landscape of the African Green Monkey Kidney-Derived Vero Cell Line. DNA Res. 21:673–683.

Patterson, N., P. Moorjani, Y. Luo, S. Mallick, N. Rohland, Y. Zhan, T. Genschoreck, T. Webster, and D. Reich. 2012. Ancient Admixture in Human History. Genetics 192:1065–1093.

Rivera-Serrano, E. E., A. S. Gizzi, J. J. Arnold, T. L. Grove, S. C. Almo, and C. E. Cameron. 2020. Viperin Reveals Its True Function. Annual Review of Virology 7:421–446.

Sène, M.-A., S. Kiesslich, H. Djambazian, J. Ragoussis, Y. Xia, and A. A. Kamen. 2021. Haplotype-resolved de novo assembly of the Vero cell line genome. npj Vaccines 6:106.

Sakuma, C., T. Sekizuka, M. Kuroda, F. Kasai, K. Saito, M. Ikeda, T. Yamaji, N. Osada, and K. Hanada. 2018. Novel endogenous simian retroviral integrations in Vero cells: implications for quality control of a human vaccine cell substrate. Scientific Reports 8:644.

Schneider, W. M., M. D. Chevillotte, and C. M. Rice. 2014. Interferon-Stimulated Genes: A Complex Web of Host Defenses. Annu. Rev. Immunol. 32:513–545.

Tang, H., and P. D. Thomas. 2016. PANTHER-PSEP: predicting disease-causing genetic variants using position-specific evolutionary preservation. Bioinformatics 32:2230–2232.

Terasima, T., M. Yasukawa, and B. Simizu. 1988. History of Vero cells in Japan. VERO scells: origin, properties and biomedical applications. Department of Microbiology School of Medicine Chiba University, Chiba, Japan:22–23.

Tukey, J. W. 1949. Comparing individual means in the analysis of variance. Biometrics:99–114.

Velikkakath, A. K. G., T. Nishimura, E. Oita, N. Ishihara, and N. Mizushima. 2012. Mammalian Atg2 proteins are essential for autophagosome formation and important for regulation of size and distribution of lipid droplets. Mol. Biol. Cell 23:896–909.

Zeng, W., L. Sun, X. Jiang, X. Chen, F. Hou, A. Adhikari, M. Xu, and Z. J. Chen. 2010. Reconstitution of the RIG-I Pathway Reveals a Signaling Role of Unanchored Polyubiquitin Chains in Innate Immunity. Cell 141:315–330.

Zhang, Y., R. Guo, S. H. Kim et al. 2021. SARS-CoV-2 hijacks folate and one-carbon metabolism for viral replication. Nat. Commun. 12:1676.

Zhao, Y. G., and H. Zhang. 2018. Autophagosome maturation: An epic journey from the ER to lysosomes. J. Cell Biol. 218:757–770.

